# Identification of a Golgi-localized peptide reveals a minimal Golgi targeting motif

**DOI:** 10.1101/2020.12.13.422530

**Authors:** Alexandra P. Navarro, Iain M. Cheeseman

**Affiliations:** Whitehead Institute for Biomedical Research, 455 Main Street, Cambridge, MA 02142; Department of Biology, Massachusetts Institute of Technology, Cambridge, MA 02142

## Abstract

Prior work has identified signal sequences and motifs that are necessary and sufficient to target proteins to specific subcellular regions and organelles such as the plasma membrane, nucleus, Endoplasmic Reticulum, and mitochondria. In contrast, minimal sequence motifs that are sufficient for Golgi localization remain largely elusive. In this work, we identified a 37 amino acid alternative open reading frame (altORF) within the mRNA of the centromere protein, CENP-R. This altORF peptide localizes specifically to the cytoplasmic surface of the Golgi apparatus. Through mutational analysis, we identify a minimal 10 amino acid sequence and a critical cysteine residue that are necessary and sufficient for Golgi localization. Pharmacological perturbations suggest that this peptide is modified through palmitoylation to promote its localization. Together, our work defines a minimal sequence that is sufficient for Golgi targeting and provide a valuable Golgi marker for live cell imaging.

## Introduction

The Golgi apparatus contributes to multiple critical cellular functions, including protein modification and sorting, vesicle transport, and lipid biosynthesis [cite a review]. The Golgi is also a highly dynamic membranous structure. Membrane trafficking through the secretory pathway results in frequent turnover of the Golgi membrane as proteins are delivered to and exported from the Golgi by vesicles (Lujan and Campelo, 2021). In addition, Golgi morphology can vary in response to cell state, such as during mitosis or apoptosis where the Golgi disassembles through a tightly-regulated processes (Liu et al., 2021). For the Golgi to perform its critical functions, resident proteins must be targeted to this organelle in a way that allows them to persist despite this reorganization. One mechanism of targeting proteins to the Golgi involves trafficking through the secretory pathway, where proteins are first targeted to the Endoplasmic Reticulum before they are transported to the Golgi (Lujan and Campelo, 2021). Once at the Golgi, proteins are retained due to their association with unique features of the Golgi such as membrane thickness or their interaction with other resident Golgi proteins (Banfield, 2011). Alternatively, cytoplasmic facing peripheral membrane proteins, such as a class of proteins called golgins, can be directly targeted to the Golgi. The association and retention of these peripheral membrane proteins at the Golgi is facilitated by lipid modifications such as myristoylation, palmitoylation, and prenylation, or indirectly through protein-protein interactions (Banfield, 2011).

For other organelles, such as nuclei or mitochondria, protein targeting requires the presence of organelle-specific signal sequences. For example, one class of nuclear localization sequence (NLS), termed ‘monopartitie’, is characterized by a cluster of 4-8 basic amino acids in which 4 or more of those amino acids are arginine or lysine residues (Lu et al., 2021). Similarly, protein targeting to the mitochondria is mediated by a signal sequence found at a protein’s N-terminus and is characterized by the presence of positively-charged and hydrophobic amino acids (Chacinska et al., 2009). In contrast, although prior work has identified protein domains that are required for the localization of resident or viral-derived Golgi proteins (Kjer-Nielsen et al., 1999a; Kjer-Nielsen et al., 1999b; Liu and Zheng, 2007; Munro and Nichols, 1999; Nozawa et al., 2003), a minimal consensus signal sequence for the directed targeting of proteins to the Golgi has not been identified.

In this work, we identify a novel 37 amino acid altORF peptide that localizes robustly to the Golgi apparatus. This altORF is expressed from the transcript of the CENP-R gene, a protein that canonically localizes to kinetochores (Hori et al., 2008). Given the unique localization behavior of this short peptide sequence, we dissected the mechanisms that underlie this localization behavior, including the palmitoylation of a critical cysteine residue and defining a minimal 8 amino acid consensus sequence. The robust localization of the altORF to the Golgi also provides a valuable imaging tool that can contribute to the analysis of dynamic Golgi behaviors. Together, this work provides valuable insight into sequences that can mediate protein targeting to the Golgi.

## Results and Discussion

### A CENPR altORF peptide localizes to the Golgi compartment

Alternative open reading frames (altORF) are sequences present in transcribed mRNAs that are distinct from the canonically-defined open reading frame. Such sequences have alternative translation start sites that result in the production of a novel protein sequence. AltORFs exist in a variety of proteins, and many have unknown functional roles and behaviors (Orr et al., 2020). In our ongoing work, W\we identified a potential altORF that initiates upstream of the defined CENP-R open reading frame and has a unique reading frame (Figure 1A). Although hypothetical, this altORF was identified based on ribosome profiling data and was included in a proteomic database of small peptides produced by altORFs with a cutoff of less than 100 amino acids (Samandi et al., 2017). This CENPR altORF corresponds to a peptide sequence of 37 amino acids with a predicted molecular weight of 4.5 kDa and a pI of 9.84. This altORF sequence is evolutionarily conserved in primates (Figure 1B, Supplemental Figure 1A), but is highly divergent or absent from other vertebrates including mice (Supplemental Figure 1B).

**Figure 1.**
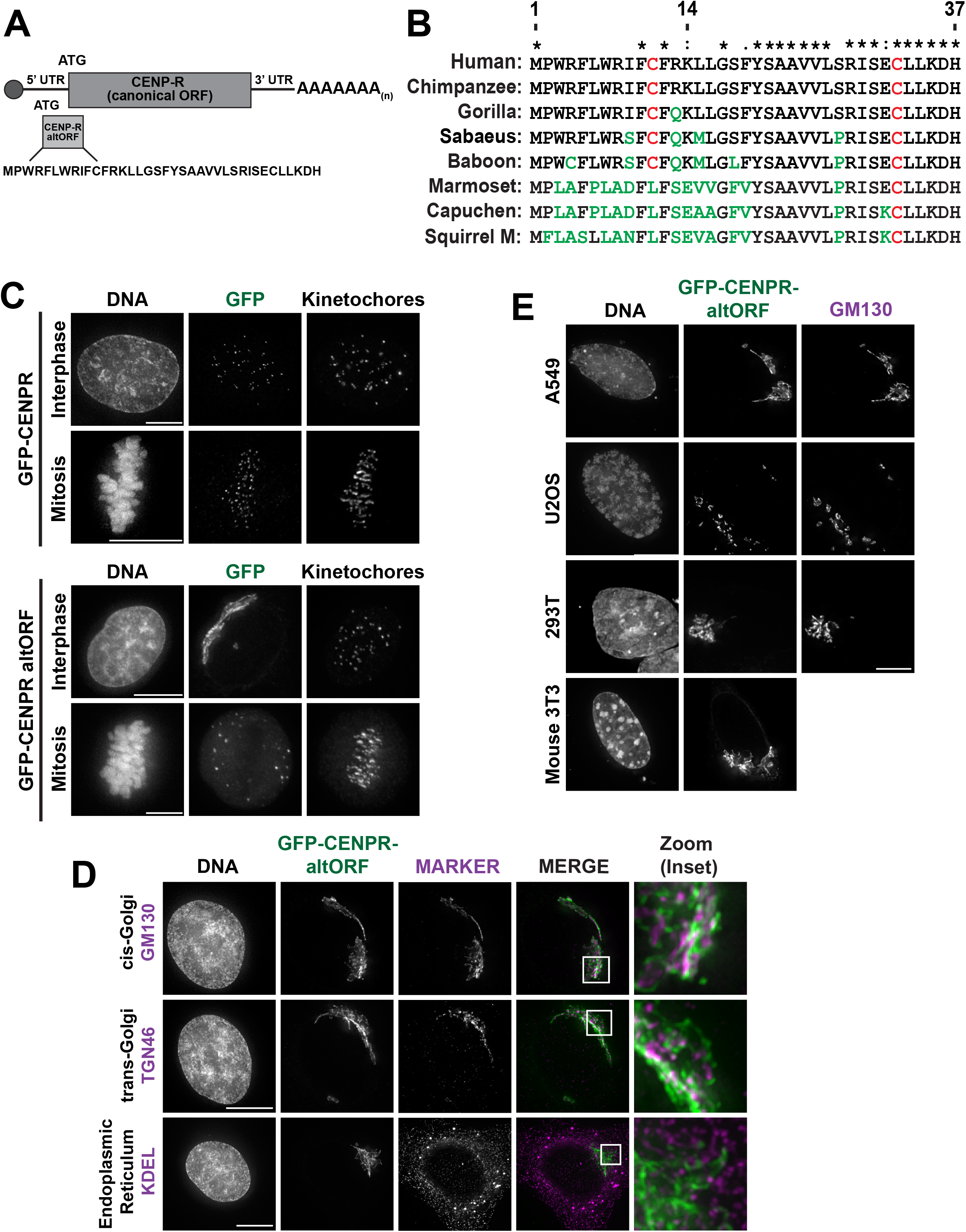
A CENP-R altORF robustly localizes to the Golgi. A. Visual representation of altORF translation from the canonical CENP-R transcript. The altORF begins upstream of the canonical ATG and has a different reading from the canonical protein. B. Multiple sequence alignment of the altORF peptide sequence across New and Old-world primates. C. Localization of CENP-R altORF during interphase and mitosis as compared to the canonical CENP-R after transfection of GFP tagged constructs in HeLa cells. The CENP-R altORF shows distinct localization compared to that of canonical CENP-R, which localizes to centromeres. GFP boost was used to amplify GFP signal and kinetochore are stained with ACA. Images in this figure are deconvolved and max projected. Scale bars, 10 μm. D. The CENP-R altORF co-localizes with the Golgi markers GM130 and TGN46, but not with the Endoplasmic reticulum marker anti-KDEL. Images were deconvolved and max projected. Scale bar, 10 μm. Inset is 5 μm. E. The CENP-R altORF localizes to the Golgi in a variety of cell lines, including mouse fibroblast 3T3 cells. The CENP-R altORF was transiently transfected into each cell lines and assessed for colocalization with Golgi marker GM130, except for mouse fibroblast as the human GM130 antibody did localize in mouse cells. Images are deconvolved and max projected. GM130 antibody recognizes the human protein so not included in 3T3 panel. Scale bar, 10 μm.

We first sought to assess the cellular behavior of this altORF peptide by ectopically expressing this altORF sequence with an N-terminal GFP tag in human HeLa cells. Strikingly, we found that this peptide displayed clear localization to what appeared to be membrane-bound structures during interphase (Figure 1C). These membranous structures localized peripherally to the nucleus and were typically compact, clustering adjacent to the nucleus. In mitosis, the altORF localized to distinct puncta throughout the cytoplasm of the dividing cell (Figure 1C). This localization behavior contrasts with that of the canonical CENP-R protein, which localizes to centromeres in both interphase and mitosis (Figure 1C). We also observed similar localization when the altORF was tagged at its C-terminus (altORF-GFP; Supplemental Figure 2A) or when expressed with a HaloTag (Supplemental Figure 2B).

**Figure 2.**
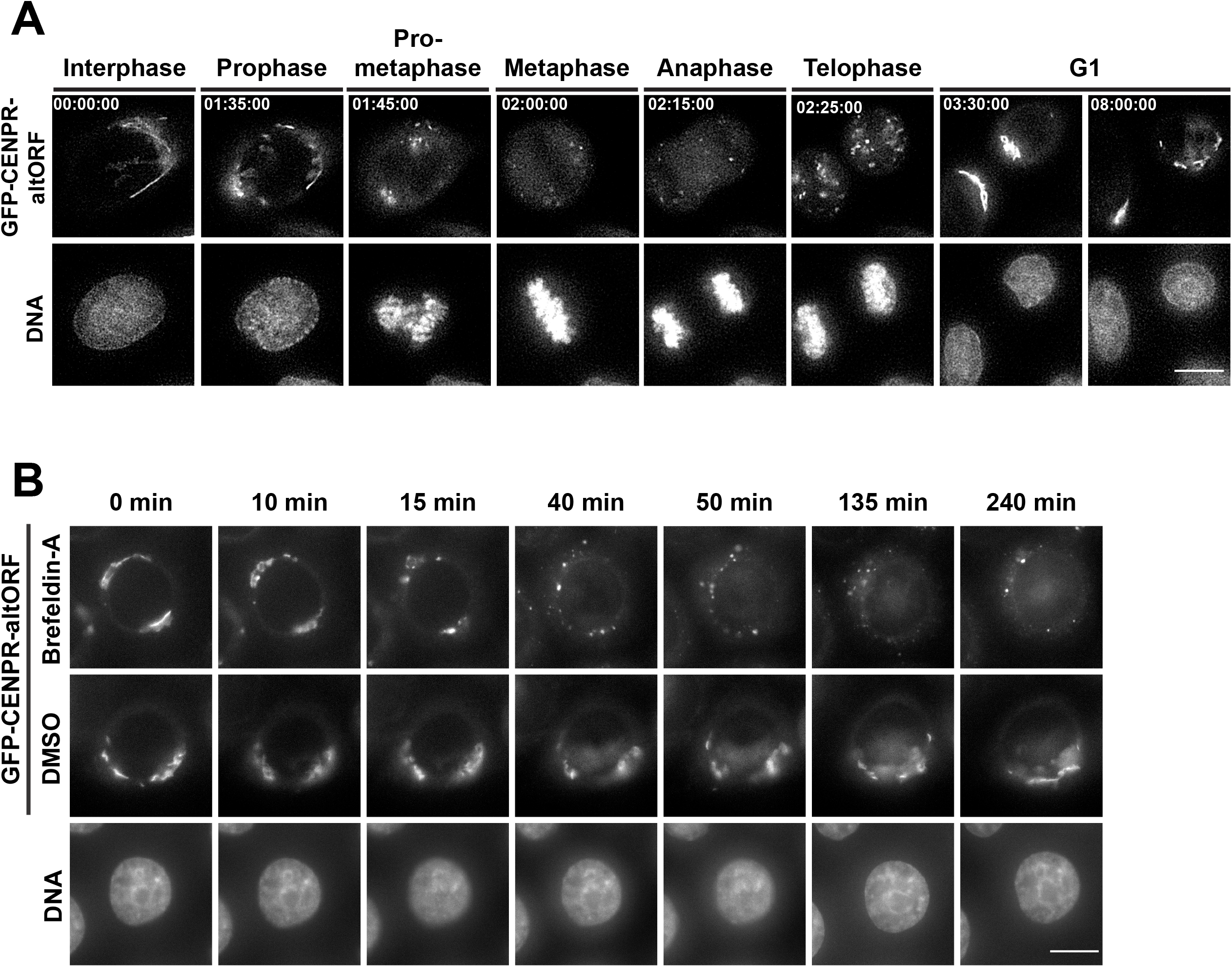
The CENP-R altORF marks the Golgi throughout dynamic processes. A. Images from a time-lapse sequence in a cell line stably expressing the CENP-R altORF. DNA was stained using sIR-DNA. Cells were imaged for 12 hours in 5-minute intervals at 37°C in CO2 independent media. The time-lapse video was deconvolved. Scale bar, 10 μm. B. Images from time-lapse sequence of a cell line stably expressing the CENP-R altORF following treatment with 0.2 μM Brefeldin A. DNA was stained with sIR-DNA. Cells were imaged immediately after the addition of Brefeldin A for 4 hours, imaged every 5 minutes. Selected frames show the breakdown of the Golgi and the association of the altORF to the Golgi throughout this process. Scale bar, 10 μm.

Based on this observed localization to a membrane compartment adjacent to the nuclear periphery, we hypothesized that the altORF likely localizes to either the Endoplasmic Reticulum (ER) or the Golgi apparatus. To test this, we compared the altORF localization with established markers for the ER – using an anti-KDEL antibody - and the Golgi – using an anti-GM130 antibody. We found that the altORF peptide did not co-localize with ER markers but overlapped closely with GM130 in fixed cells (Figure 1D). The Golgi network is further divided into cis-, medial-, and trans-compartments. To define the localization behavior of this altORF peptide, we additionally compared its localization to that of GM130, a marker of the cis-Golgi, and TGN46, a marker of the trans-Golgi (Kain et al., 1998; Nakamura et al., 1995). In each case, we observed approximate co-localization of the altORF peptide with these antibodies, but the resolution limits of these images did not allow us to precisely define which sub-compartment within the Golgi is labeled by this peptide (Figure 1D). Together, we conclude that this altORF peptide localizes to Golgi apparatus in HeLa cells.

We next sought to test whether the observed localization behavior for the CENPR altORF also occurred in other cell lines. To test this, we transfected the N-terminal GFP construct into a panel of human cell lines including the lung carcinoma A549 cell line, the osteosarcoma U2OS cell line, and the human embryonic kidney 293T cell line. In each case, we observed similar localization to the nuclear periphery as in HeLa cells, as well as co-localization with GM130 (Figure 1E). In addition to these human cell lines, we tested the localization of the human altORF in mouse fibroblast NIH3T3 cells. Although this altORF sequence is absent from mice, we observed a similar localization of the altORF to the Golgi in NIH3T3 cells (Figure 1E). We therefore conclude that the altORF robustly localizes to the Golgi across a variety of mammalian cell lines.

### The CENPR altORF remains associated during Golgi remodeling and dynamics

To assess the dynamic localization of the altORF to the Golgi, we used the altORF construct to generate a cell line stably expressing the N-terminally tagged GFP-altORF using retroviral integration. Stable expression of the altORF had no apparent effect on cell growth (Supplemental Figure 2C). Using this cell line, we analyzed the association of the altORF with the Golgi under conditions where Golgi morphology is dynamic. First, we visualized the localization of the altORF peptide throughout the cell cycle. The Golgi undergoes a cycle of assembly and disassembly as cells progress through cell division (Tang and Wang, 2013). Using time lapse imaging, we observed the rapid fragmentation of the Golgi upon mitotic entry, with the altORF appearing as small puncta throughout the cytoplasm of the dividing cell, consistent with prior fixed cell analysis. As cells exited mitosis in telophase, we observed the reformation of larger stacks and ultimately the full reformation of the Golgi into ribbon-like structures in G1 (Figure 2A).

Next, we sought to visualize altORF localization and Golgi behavior by inducing morphological changes to the Golgi using Brefeldin A treatment. Brefeldin A is an inhibitor of the secretory pathway that causes the Golgi to disassemble and redistribute into the Endoplasmic reticulum (Fujiwara et al., 1988). Upon treatment with Brefeldin A, we observed the initiation of the breakdown of the ribbon-like structures of the Golgi into small puncta that moved freely throughout the cytoplasm of the cell within 15 minutes of treatment (Figure 2B). Thus, the altORF peptide remains robustly associated with the Golgi membrane under conditions where the organelle is disassembled, providing a valuable tool for the live cell imaging of Golgi dynamics.

### The CENPR altORF localizes to the cytoplasmic surface of the Golgi

The robust localization of this 37 amino acid altORF peptide to the Golgi raises questions as to how this peptide associates with and is retained at the Golgi. We first sought to determine whether this altORF peptide is associated with the cytoplasmic or luminal side of the Golgi membrane. To test this, we utilized a linker that contains a TEV protease cleavage site between GFP and the start of the altORF sequence (Cheeseman and Desai, 2005). We then expressed mCherry-tagged TEV protease in cells stably expressing this GFP-TEV-altORF peptide fusion. If the altORF peptide faces the cytoplasm, then the TEV cleavage site should be exposed and susceptible to cleavage by the cytoplasmic TEV protease, thereby resulting in the loss of Golgi-localized GFP signals. Indeed, upon TEV expression, we observed a substantial increase in cytoplasmic GFP fluorescence and a loss of Golgi-specific GFP localization (Figure 3A), suggesting that the altORF localizes to the cytoplasmic surface of the Golgi.

**Figure 3.**
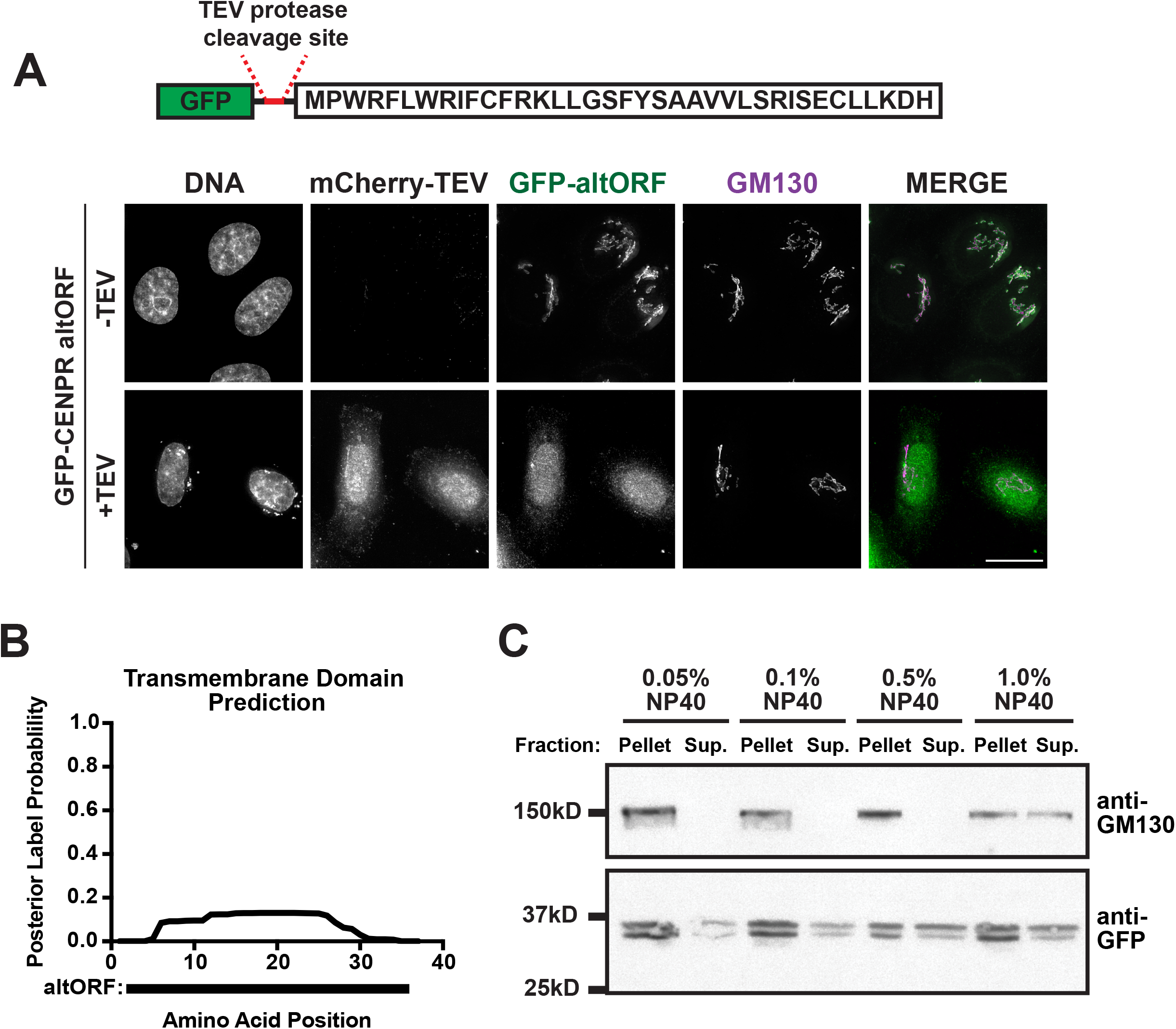
CENP-R altORF associates to the cytoplasmic surface of the Golgi. A. Localization of the GFP-CENPR altORF peptide stably expressed in HeLa cells in the presence of transfected mCHerry-TEV protease construct. Localization to the Golgi was determined based on colocalization with Golgi marker GM130. Images are deconvolved and maximally projected. Scale bar, 10 μm. B. Hydrophobicity plot of the CENPR altORF peptide sequence. Sequence was run through the Phobius program (https://phobius.sbc.su.se/poly.html), probability values assigned to amino acids to determine transmembrane topology are plotted. C. Western showing the presence of GFP-altORF or GM130 following lysis with buffers of various amounts of the detergent NP40.

We next sought to elucidate the sequence requirements for targeting the altORF peptide to the Golgi. Resident Golgi proteins can associate with the Golgi membrane as transmembrane or peripheral membrane proteins. Golgi-associated transmembrane domains typically span ~20 aa in length and are enriched in aromatic and hydrophobic amino acids (Sharpe et al., 2010). To determine whether this altORF peptide could act as a transmembrane protein at the Golgi, we utilized the Phobius online transmembrane topology tool to test for the presence of predicted transmembrane domains (Kall et al., 2004; Kall et al., 2007). Analysis of the altORF sequence alone found a low probability for the presence of a transmembrane domain, suggesting that the altORF likely associates with the Golgi as a peripheral membrane protein (Figure 3B). To further assess the association of the altORF with the Golgi, we used cell lysis in the presence of NP40 to assess the conditions required to solubilize the altORF peptide. We compared this to the solubilization behavior to that of GM130, a well-characterized Golgi protein known to act as a strongly-associated peripheral membrane protein (Nakamura et al., 1995). We found that the altORF peptide was solubilized in the presence of 0.05% NP40 detergent whereas GM130 required 1% NP40 to be liberated to the supernatant fraction (Figure 3C). This indicates that the altORF is more weakly associated with the Golgi relative to GM130.

Some proteins that are peripherally associated with the Golgi do so through protein-protein interactions with Golgi-resident proteins. To determine whether altORF association with the Golgi was mediated through its interaction with another Golgi-associated protein, we sought to identify altORF-interacting proteins by immunoprecipitation-mass spectrometry (IP-MS) analysis. Although we were able to isolate the altORF peptide as a GFP fusion, we did not detect any additional Golgi-associated proteins in our mass spec analysis relative to controls (data not shown). Thus, although our IP experiments do not exclude the possibility that protein-protein interactions help to promote altORF localization, robust and persistent protein-protein interactions are unlikely to be responsible for retaining the altORF at the Golgi.

### Cysteine residues are important for localization of the altORF peptide to the Golgi

We next sought to determine the amino acid residues that are required for Golgi localization. The altORF peptide has regions with hydrophobic amino acid residues and aromatic residues, and contains two cysteine residues (Supplemental Figure 2D), which may play roles in Golgi localization (Banfield, 2011). To test the contributions of these residues, we generated truncation mutants to determine the minimal sequence required for Golgi localization (Figure 4A). We found that truncating the last 23 amino acids did not affect Golgi localization as the GFP-altORF^21aa^ and the GFP-altORF^14aa^ mutants colocalized with GM130 (Figure 4B). However, truncating the altORF peptide sequence down further to its first 9 amino acids resulted in the loss of Golgi localization and an increase in the cytoplasmic GFP signal (Figure 4B). Next, we truncated an additional 4 amino acids from the N-terminus of the altORF^14aa^ mutant resulting in a 10 amino acid sequence (amino acids 5-14). This 10 amino acid sequence was sufficient to direct Golgi localization (Figure 4B). Of the residues eliminated through the truncation of the altORF sequence from 14 to 9 amino acids, the loss of the Cys11 residue was particularly interesting given its potential to be modified (Figure 4A). To test the role of cysteine residues in altORF localization, we generated an altORF mutant in which both cysteine residues were mutated to alanine (altORF^C11A, C32A^). The altORF^C11A, C32A^ mutant failed to localize robustly to the Golgi (Figure 4C), suggesting that the cysteine residues play a critical role in targeting and retaining this peptide to the Golgi.

**Figure 4.**
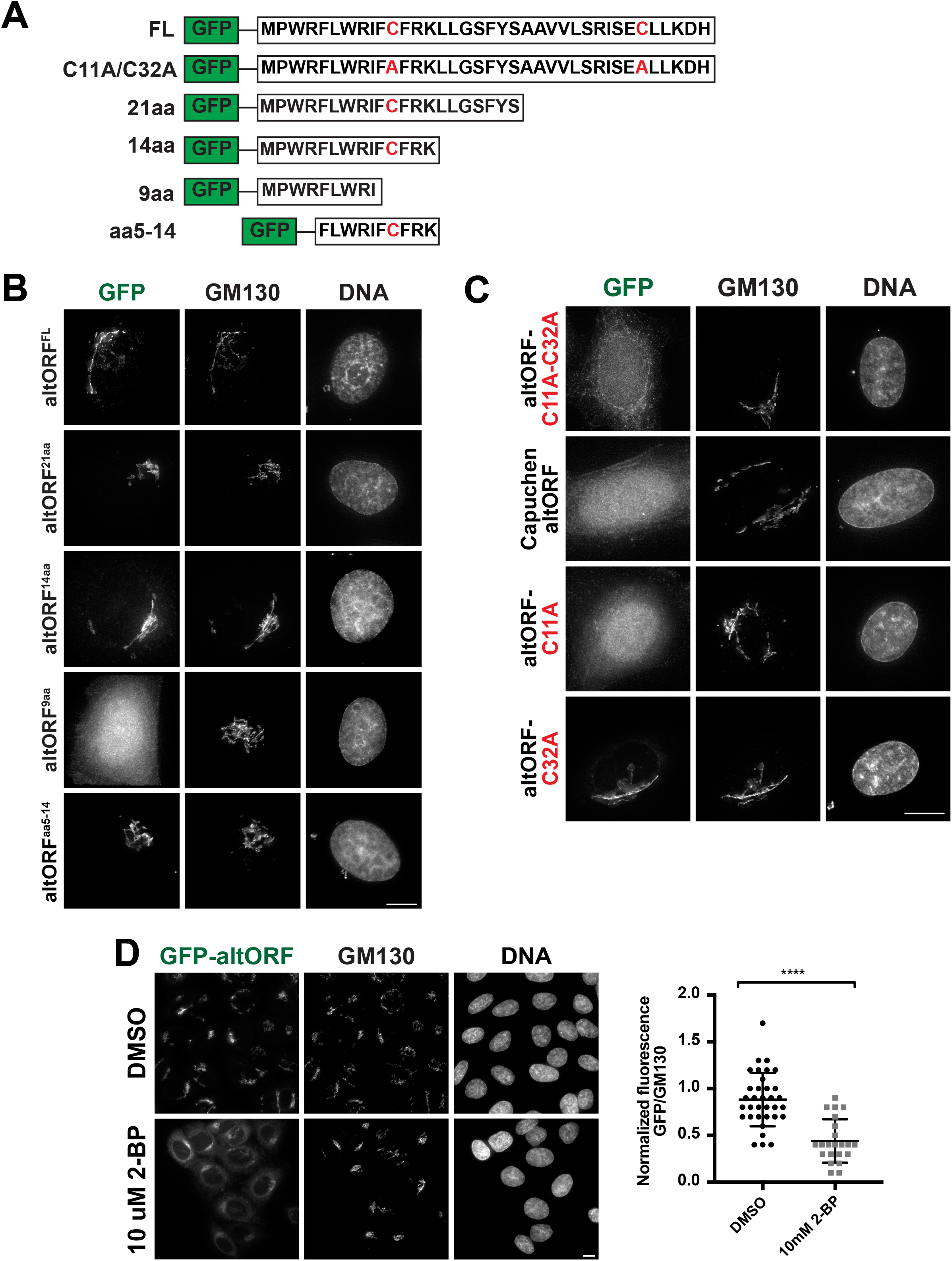
Cysteine residues are required for altORF localization to the Golgi. A. Figures representing the truncation mutants generated and tested for localization. B. Localization of the truncated altORF constructs transfected in HeLa cells. Localization to the Golgi was determined based on colocalization with Golgi marker GM130. Cells were fixed and imaged 48 hours post transfection. Images are deconvolved and max projected. Scale bar, 10 μm. C. Localization of the dual cysteine mutant altORF constructs transfected in HeLa cells. Localization to the Golgi was determined based on colocalization with Golgi marker GM130. Cells were fixed and imaged 48 hours post transfection. Images are deconvolved and max projected. Scale bar is 10 μm. D. Localization of the altORF peptide following treatment with 2-Bromopalmitate. Cells were treated with either DMSO or 10 uM 2-Bromopalmitate for 3 hours before fixation. Images are deconvolved. Scale bar is 10 μm. GFP signal is graphed as a ratio of GM130 fluorescence per cell. Quantification of GFP signal in DMSO and 2-BP treated cells with N=34 and N=22, respectively. Error bars represent the mean and standard deviation. Statistical significance **** p <0.0001 as determine by Mann-Whitney test.

We next took an evolutionary-guided approach to define the requirements for altORF Golgi localization. The altORF sequence present in New World primates differs from that in Hominids and Old-World primates due to an insertion and deletion event which disturb and restore the reading frame resulting in a peptide sequence that differs from the human altORF peptide sequence (Supplemental Figure 1A, 1B; Harmit Malik, personal communication). Although both the C11 and C32 cysteine residues are present in the Old-World primate altORF peptide sequences, only the C32 residue is conserved in New World primates (Figure 1B). To evaluate the consequences of these changes in Golgi localization, we tested the localization of the Capuchin altORF sequence, which lacks the C11 residue. Using transfection of a GFP-tagged construct into HeLa cells, we found that the Capuchin altORF sequence did not localize to the Golgi (Figure 4C).

Based on both our observations from the mutational analysis and the evolutionary conservation of the altORF sequence, we hypothesized that the Cys11 residue would be required for Golgi localization. To test this, we generated altORF mutants in which individual cysteine residues were mutated to alanine (altORF^C11A^ and altORF^C32A^). We found that the altORF^C11A^ failed to localize to the Golgi and was instead localized diffusely throughout the cytoplasm (Figure 4C). In contrast, the altORF^C32A^ mutant behaved similarly to the wild type sequence localizing to the Golgi (Figure 4C). We therefore conclude that the Cys11 residue is necessary for the altORF to localize to the Golgi.

### Lipid modification of the altORF peptide contributes to Golgi localization

Our results suggest that the altORF peptide associates with the Golgi as a peripheral membrane protein and that the cysteine residues are required for this localization. Therefore, we hypothesized that the localization of the altORF to the Golgi is mediated by a post-translational modification that targets the cysteine residues in this peptide. Potential post-translational modifications that occur at cysteine residues and are known to contribute to Golgi localization include myristoylation, prenylation, and palmitoylation (Aicart-Ramos et al., 2011; Banfield, 2011; Palsuledesai and Distefano, 2015; Udenwobele et al., 2017). Due to the absence of clear myristoylation or prenylation consensus motifs within the altORF sequence (Palsuledesai and Distefano, 2015; Udenwobele et al., 2017), we hypothesized that palmitoylation may be involved in facilitating the localization of the altORF to the Golgi. Palmitoylation is a reversible post-translational lipid modification that typically functions to enhance protein hydrophobicity and promote the membrane association of modified proteins (Guan and Fierke, 2011). The process of palmitoylation is catalyzed by a large family of enzymes termed palmitoyl acyltransferases (PATs), consisting of 23 distinct proteins that all share a core DHHC motif (Philippe and Jenkins, 2019). Most of these enzymes are enriched at the Golgi, although they additionally localize to the ER and plasma membrane (Ernst et al., 2018; Ohno et al., 2006; Rocks et al., 2010). The function of these PAT enzymes is largely redundant, as multiple PAT enzymes have been shown to have overlapping substrates. Currently, no generalized consensus motifs have been identified for palmitoylated proteins (Salaun et al., 2010).

To test whether palmitoylation is involved in altORF Golgi localization, we first sought to identify PAT enzymes that are required for altORF localization. Utilizing, a CRISPR/Cas9 inducible knockout strategy (McKinley and Cheeseman, 2017), we specifically targeted the PAT enzymes that localize to the Golgi, including ZDHHC3, ZDHHC7, ZDHHC9, ZDHHC11, ZDHHC13, ZDHHC15, ZDHHC17, ZDHHC21, and ZDHHC22 (Ernst et al., 2018; Rocks et al., 2010). However, likely due to the functional redundancy of these enzymes, knocking out individual enzymes or two enzymes combinations (specifically ZDHHC 3/7; 3/9, 7/9) did not result in the mis-localization of the altORF (data not shown) (Greaves et al., 2017; Philippe and Jenkins, 2019; Rocks et al., 2010). Thus, as an alternative approach, we sought to pharmacologically perturb cellular palmitoylation utilizing the inhibitor 2-Bromopalmitate (2-BP), a palmitate analog that inhibits this process (Salaun et al., 2010). Following treatment with 2-BP for 4 hours, we observed the mislocalization of the altORF from the Golgi and an increase in cytoplasmic GFP signal (Figure 4D). This observation supports a model in which the Golgi localization of the altORF relies on active palmitoylation. Together, our work suggests that cysteine 11 is modified by palmitoylation to promote the localization of the altORF to the Golgi.

### Determination of a minimal Golgi-targeting sequence

Given the critical role the Cys11 residue plays in the localization of this peptide, we next sought to determine what other amino acid residues within the minimal 10 amino acid sequence, identified in the altORF^aa5-14^ truncation mutant, are required for Golgi localization (Figure 4A). Within this 10-amino acid sequence, there is a ‘FCF’ motif that is conserved across Old World primates with varying amino acids surrounding this motif (Figure 1B). Therefore, we sought to assess the functional importance of the amino acids directly surrounding the cysteine residue. Mutating the phenylalanine residues to tyrosine to retain their aromatic character resulted in a loss of localization. In contrast, mutating these residues to leucine to preserve the hydrophobicity did not affect Golgi localization (Figure 5A). This suggests that the presence of hydrophobic amino acids on either side of the cysteine is important for Golgi localization.

**Figure 5.**
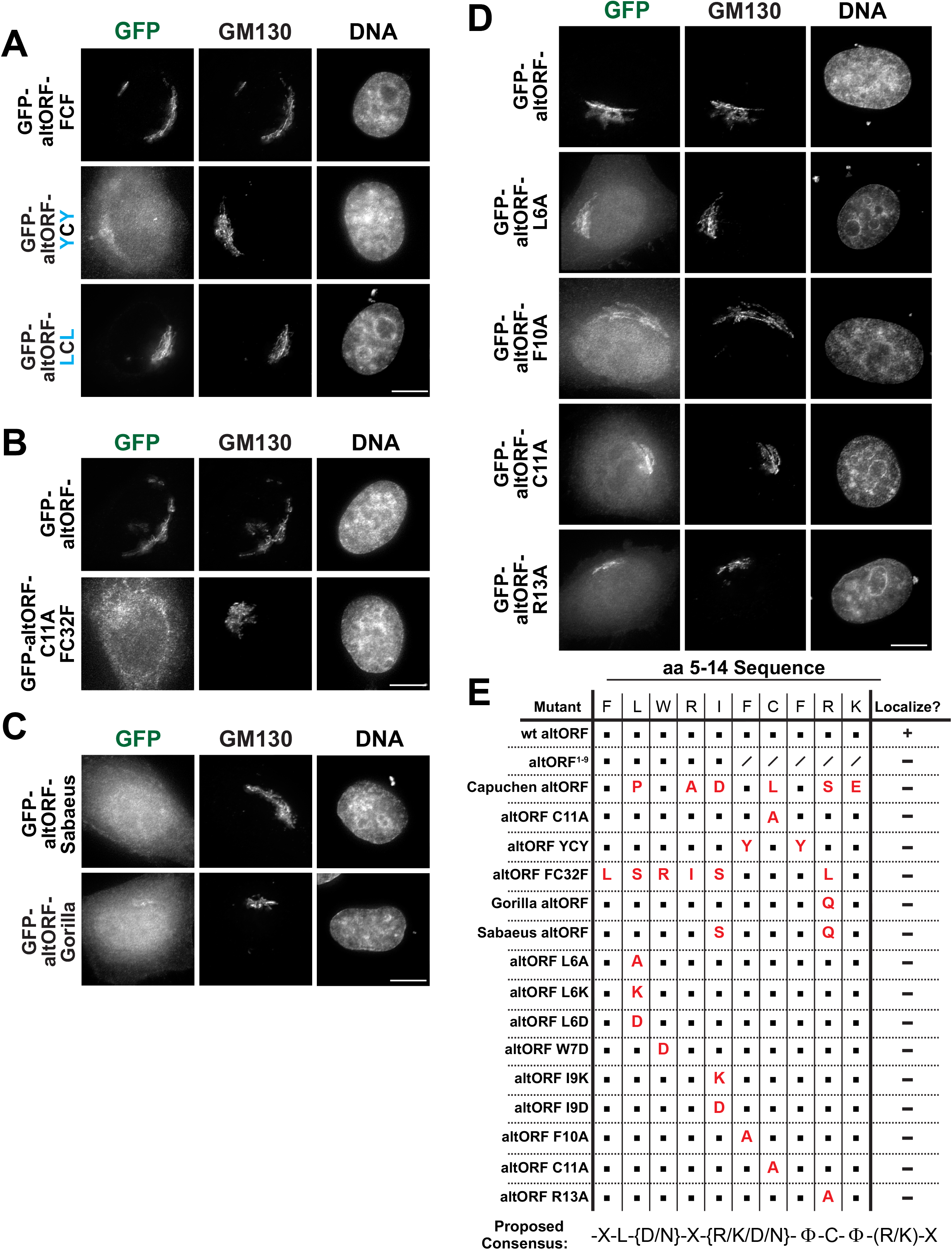
FCF motif not sufficient to confer Golgi localization. A. Representative images of the FCF mutant localization in HeLa cells. Cells were fixed and imaged 48 hours post transfection. Images are deconvolved and max projected. Scale bar is 10 μm. B. Localization of GFP tagged altORF mutant testing sufficiency of FCF motif. Constructs were transiently transfected in HeLa cells and expressed for 48 hours prior to fixation. Images are deconvolved and max projected. Scale bar, 10 μm. C. Localization of GFP tagged Sabaeus altORF and motif mutant testing sufficiency of FCF motif. Constructs were transiently transfected in HeLa cells and expressed for 48 hours prior to fixation. Images are deconvolved and max projected. Scale bar, 10 μm. D. Representative images of the Alanine mutants of the 10 amino acid peptide sequence. Constructs were transiently expressed in HeLa cells for 48 hours prior to fixation. Images were deconvolved and max projected. Scale bar, 10 μm. E. Table summarizing the mutants that failed to localize and the proposed consensus sequence required for Golgi localization. In the consensus sequence the () brackets represent the amino acids that can be substituted into that position; and the {} brackets represent amino acids that cannot be included within a given position.

Next, we examined whether the “FCF” motif is sufficient to direct Golgi localization. To test this, we took the altORF^C11A^ mutant, which does not localize to the Golgi (Figure 4B) and mutated the two amino acid residues adjacent to the C32 residue to phenylalanine (Supplemental Figure 2D). This mutant failed to localize to the Golgi indicating that the ‘FCF’ motif is not sufficient to confer Golgi localization (Figure 5B). Finally, we tested the functional importance of the amino acid residues surrounding the ‘FCF’ motif. First, we utilized the evolutionarily conserved altORF sequences to probe whether the surrounding amino acids are required to promote the Golgi localization. Based on this analysis, we selected the altORF sequence from two distinct primates - the Gorilla sequence that contains one amino acid substitution, and the Sabaeus sequence that contains two amino acid substitutions. Despite these limited changes, neither sequence localized to the Golgi (Figure 5B) suggesting stringent sequence requirements for Golgi localization. Thus, we took an unbiased approach to dissect the amino acid requirements at each position within the altORF sequence. To do this, we systematically performed site directed mutagenesis in which we targeted each amino acid in the peptide sequence (Figure 5D; Supp Fig 3A). We generated mutant constructs in which we individually mutated each amino acid within the peptide sequence to alanine or an amino that alters the characteristics of a given residue to determine which amino acids can be substituted at each position (Supp Fig 3A). From this analysis, we established the following generalized motif required for Golgi localization of this peptide: X-[ILVM]-{DN}-X-{RKDN}-[FILVM]-C-[IFLVM]-X-[RK]-X (Figure 5E).

In conclusion, our work identifies a small, 37 aa peptide derived from the CENPR transcript that localizes robustly to the Golgi apparatus. This localization behavior is dependent on the Cys11 residue present within the peptide. Chemical perturbation of cellular palmitoylation disrupted Golgi localization of the altORF suggesting that palmitoylation is likely the post-translational modification responsible for its Golgi localization. We further identify a minimal 10 amino acid sequence that is sufficient to target GFP to the Golgi, suggesting that this sequence could function as a Golgi-targeting sequence mediated by palmitoylation. In addition to revealing the requirements for Golgi localization and palmitoylation, this altORF peptide provides a valuable marker for visualizing the Golgi apparatus in intact cells. This small peptide, when tagged with a fluorescent protein, can be used to image the Golgi by either transient transfection or stable expression, and remains associated with the Golgi throughout its dynamic rearrangement. The altORF can also be used in a variety of cell lines, including mouse fibroblasts, for both fixed and live imaging of the Golgi. We hope that this easy-to-use Golgi labeling construct will provide a valuable tool for researchers conducting studies of Golgi localization and dynamics in addition to the existing Golgi labeling proteins (Aicart-Ramos et al., 2011; Kall et al., 2004; Udenwobele et al., 2017).

## Acknowledgements

The authors thank the members of the Cheeseman lab for their support, and Lawrence Welch, Isheir Raote, Yamuna Krishnan, Sam Campos, and John Ngo for their suggestions and comments (via Twitter). Additional thanks to Harmit Malik for his help in analyzing the evolutionary conservation of this altORF sequence. This work was supported by grants from the National Science Foundation (2029868) and the NIH/National Institute of General Medical Sciences (R35GM126930) to IMC and a National Science Foundation Graduate Research Fellowship to A.P.N.

## Methods

### Cell Culture, cell transfection, and cell line generation

HeLa, U2Os, A549, and 3T3 cells were cultured in Dulbecco’s Modified Eagle Medium supplemented with 10% fetal bovine serum (GE Healthcare), 100 U/ml penicillin and streptomycin, and 2 mM L-glutamine at 37C with 5% CO2. Stable cell line expressing altORF was generated by retroviral infection followed by selection with Blasticidin at 2 ug/mL and single cell sorting to generate a clonal cell line. Transfection was performed with Xtremegene-9 as per product protocol.

### Immunoblotting

Cells were harvested from cell lines stably expressing the GFP-altORF. Cells were harvested by trypsinization with 0.05% Trypsin/EDTA and re-suspended in DMEM. Cells were counted and pellets with 2 million cells were collected. Cells were then lysed in lysis buffer (50 mM HEPES, pH 7.4; 1 mM EGTA; 1mM MgCl2; 100 mM KCl; 10% glycerol; 1X Complete EDTA-free protease inhibitor cocktail (Roche), 1 mM PMSF, 20 mM beta-glycerophosphate, 1mM sodium fluoride, and 0.4 mM sodium orthovanadate, pH 7.4) with the corresponding percentage of NP40. Lysates were then sonicated. Cellular debris was removed by centrifugation. Samples were separated by SDS-PAGE and transferred to PVDF membrane. Membranes were blocked for 1 hour in Blocking buffer (2% nonfat dry milk + 0.1% Tween-20 in TBS). Primary antibodies (anti-GFP from Roche Prod No. 1181446001; anti-GM130 from Cell Signaling D6B1 #12480) were diluted in 0.2% nonfat dry milk in TBS +0.1% Tween and incubated on membrane overnight at 4C. HRP-conjugated secondary antibodies (antiMouse-HRP from Biorad #1705047; antiRabbit-HRP from Kindle Biosciences #R1006) were diluted 1:10,000 in 0.2% nonfat dry milk in TBS +0.1% Tween for 1 hour at room temperature. Clarity enhanced chemiluminescence substrate (Bio-Rad) was added to the membrane according to the manufacturer’s instructions. Membranes were imaged with a KwikQuant Imager (Kindle Biosciences).

### Immunofluorescence and Microscopy

Cells were plated on glass coverslips coated with poly-L-lysine (Sigma-Aldrich). Cells were fixed with 4% formaldehyde in PBS for 10 min at room temperature or with Methanol for 5 minutes at −20C. Blocking and all antibody dilutions were performed using Abdil (20 mM Tris, 150 mM NaCl, 0.1% Triton X-100, 3% BSA and 0.1% NaN3, pH 7.5). PBS plus 0.1% Triton X-100 (PBS-TX) was used for washes. GFP-Booster from Chromotek was used to amplify the fluorescence of the GFP-tagged transgenes at a dilution of 1:200. Microtubules were stained with DM1A antibody from Sigma-Aldrich at a dilution of 1:3000. For staining of Golgi for co-localization experiments GM130 antibody from Cell Signaling Technologies (#12480S) at a dilution of 1:3200 and TGN146 antibody from Abcam (ab50595) at a dilution of 1:500. For staining of the Endoplasmic Reticulum KDEL antibody from abcam (ab176333) was used at a dilution of 1:100. Cy3- and Cy5-conjugated secondary antibodies (Jackson ImmunoResearch Laboratories) were used at a 1:200 dilution in PBS plus 0.1% Triton X-100. DNA was visualized by incubating cells in 1 ug/ml Hoechst33342 (Sigma-Aldrich) in PBS plus 0.1% Triton X-100 for 10 min. Coverslips were mounted using PPDM (0.5% p-phenylenediamine and 20 mM Tris-Cl, pH 8.8, in 90% glycerol) and sealed with nail polish. For live-cell imaging, cells were seeded into 8-well glass-bottomed chambers (Ibidi) and moved into CO2-independent media (LifeTech) before imaging at 37°C. For certain movies, DNA was stained with Images were acquired on a DeltaVision Core deconvolution microscope (Applied Precision) equipped with a CoolSnap HQ2 charge-coupled device camera (Photometrics). Images were maximally projected and deconvolved where noted.

### Small Molecule Treatments

Cells were treated with 0.2uM Brefeldin A (Sigma B5936) resuspended in DMSO. 2-Bromopalmitate (Sigma 21604) was resuspended in DMSO and used at 10 uM. Cells were treated for 4 hours before fixation.

## Supplemental figure legends

**Supplemental Figure1. Conservation and alignment of altORF sequence across multiple organisms.**

A. Multiple sequence alignment of altORF sequences across New and Old-world primates.

B. Global alignment of the altORF sequence between mice and humans.

**Supplemental Figure 2. altORF localization is not affected by C-terminal tagging or use of alternative tag.**

A. The altORF peptide localizes to the Golgi whether it is tagged at the N- or C-terminus. Constructs were transiently transfected into HeLa cells and assessed for colocalization with Golgi marker GM130. Images are deconvolved and maximally projected. Scale bar, 10 μm.

B. Localization of Halo tagged altORF sequence and colocalization with GM130. Construct was transfected into HeLa cells. HaloTag conjugated to JF646 dye. Images are deconvolved and mac projected. Scale bar, 10 μm.

C. Total cell counts were determined for Hela and the cell line stably expressing GFP-altORF over the course of 6 days. Counts were repeated a total of three times.

D. Table representing the altORF peptide highlighting the amino acids belonging to each group of interest.

**Supplemental Figure 3. Summary of altORF mutants generated and their localization behaviors.**

A. Table summarizing all altORF mutants generated and their localization behaviors. A (+) indicates robust Golgi localization, (~) indicates robust golgi localization with some cytoplasmic signal, and (−) indicates either complete mislocalization of the peptide or some Golgi localization with a lot of cytoplasmic signal. In the consensus sequence, as noted, the () brackets represent the amino acids that can be substituted into that position; and the {} brackets represent amino acids that cannot be included within a given position.

## Notes

### Competing Interest Statement

The authors have declared no competing interest.

### Summary of Updates

This revised version includes new experiments to analyze the sequence requirements for the localization of the CENPR altORF to the Golgi.

