## Supplemental Figure 1 for "Identification of a Golgi-localized peptide reveals a minimal Golgi targeting motif"

## A

### Alignment of Old World Primates/Hominoids altORF sequence

```
Human    ATGCCTTGGCGTTTCCTTTGGCGGATTTTCTGTTTTCGGAAGTTGCTGGGTTCGTTTTATTCAGCGGCAGTGGTGCTTTCCCGAATCTCAGAATGCCTGTTAAAAGATCACTGA
Chimp    ATGCCTTGGCGTTTCCTTTGGCGGATTTTCTGTTTTCGGAAGTTGCTGGGTTCGTTTTATTCAGCGGCAGTGGTGCTTTCCCGAATCTCAGAATGCCTGTTAAAAGATCACTGA
Gorilla  ATGCCTTGGCGTTTCCTTTGGCGGATTTTCTGTTTTCAGAAGTTGCTGGGTTCGTTTTATTCAGCGGCAGTGGTGCTTTACCGAATCTCAGAATGCCTGTTAAAAGATCACTGA
Baboon   ATGCCTTGGTGTTTCCTTTGGCGGAGTTTCTGTTTTCAGAAGATGCTGGGTTTGTTTTATTCAGCGGCAGTGGTGCTTCCCCGAATCTCAGAATGCCTGTTAAAAGATCACTGA
```

### Alignment of New World Primates altORF sequence

```
Human            ATGCCT-TGGCGTTTCCTTTGGCGGATTTTCTGTTTTCGGAAGTTGCTGGGTTCGTTTTATTCAGCGGCAGTGGTGCTTTCCCGAATCTCAGAATGCCTGTTAAAAGATCACTGA
Capuchen         ATGCCTCTGGCGTTTCCTTTGGCGGATTTTCTGTTCTCAGAAGCTGCTGGGTTCG-TTTATTCAGCCGCAGTGGTGCTTCCCCGAATCTCAAAATGCCTGTTAAAAGATCACTGA
Marmoset         ATGCCTCTGGCGTTTCCTTTGGCGGATTTTCTGTTCTCAGAAGTTGTTGGGTTCG-TTTATTCAGCGGCAGTGGTGCTTCCCCGAATCTCAGAATGCCTGTTAAAAGATCACTGA
Squirrel Monkey  ATGTTTCTGGCGTCTCTTTTGGCGAATTTTCTGTTCTCAGAAGTTGCTGGGTTCG-TTTATTCAGCGGCAGTGGTGCTTCCCCGAATCTCAAAATGCCTGTTAAAAGATCACTGA
```

## B

### Alignment of Mammalian altORF sequence

```
         M P W R F  L W R  I  F  C  F  R  K  L  L  G  S  F  Y  S  A  A  V  V  L  S  R  I  S  E  C  L  L  K  D H  *
Human    ATGCCTTGGCGTTTCCTTTGGCGGATTTTCTGTTTTCGGAAGTTGCTGGGTTCGTTTTATTCAGCGGCAGTGGTGCTTTCCCGAATCTCAGAATGCCTGTTAAAAGATCACTGA
         **** *  ......        ....   .....          .....          ... .  . ....  . . .      ......         ...... .....
Mouse    ATGCTTCCGCGTTT------GCGG--TTTCT--------CAGTTG--------GTTCTGTCCAGCAACGGAG------TCCCGA-------AATGCCAGTTAA-----------
         M L P R    L     R  F     L    S     W     F C P A T E         S R          N A S *
```
