## Supplemental Figure 2 for "Identification of a Golgi-localized peptide reveals a minimal Golgi targeting motif"

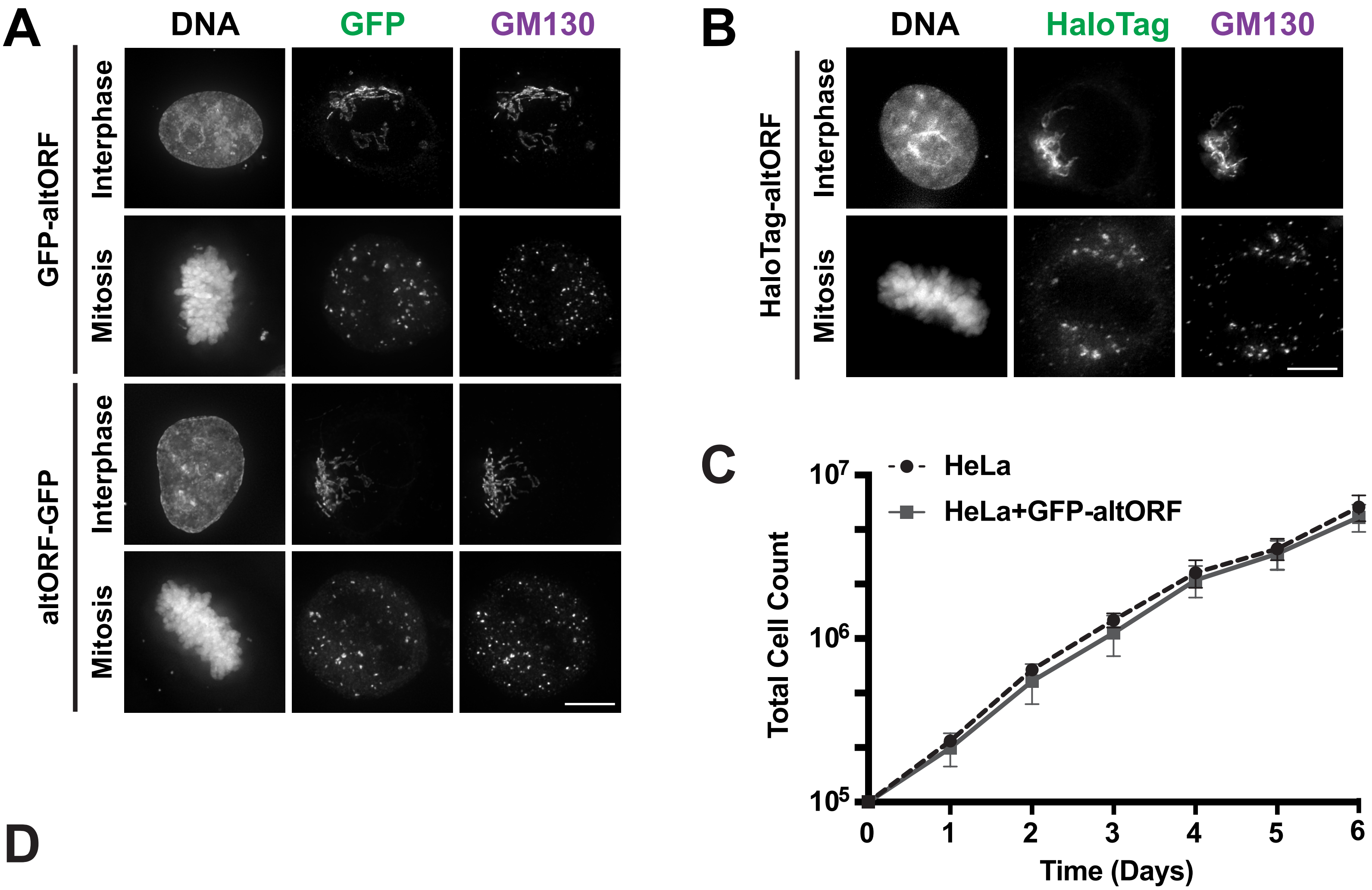


**D**

| | Sequence |
|---|---|
| Hydrophobic aa | MPWRFLWRIFCFRKLLGSFYSAAVVLSRISECLLKDH |
| Aromatic aa | MPWRFLWRIFCFRKLLGSFYSAAVVLSRISECLLKDH |
| Cysteine PTM | MPWRFLWRIFCFRKLLGSFYSAAVVLSRISECLLKDH |
