## Supplemental Figure 3 for "Identification of a Golgi-localized peptide reveals a minimal Golgi targeting motif"

## A

| Mutant | F | L | W | R | I | F | C | F | R | K | Localize? |
|---|---|---|---|---|---|---|---|---|---|---|---|
| | **aa 5-14 Sequence** | | | | | | | | | | |
| wt altORF | ■ | ■ | ■ | ■ | ■ | ■ | ■ | ■ | ■ | ■ | + |
| altORF YCY | ■ | ■ | ■ | ■ | ■ | Y | ■ | Y | ■ | ■ | — |
| altORF LCL | ■ | ■ | ■ | ■ | ■ | L | ■ | L | ■ | ■ | + |
| altORF F5A | A | ■ | ■ | ■ | ■ | ■ | ■ | ■ | ■ | ■ | + |
| altORF F5K | K | ■ | ■ | ■ | ■ | ■ | ■ | ■ | ■ | ■ | + |
| altORF F5D | D | ■ | ■ | ■ | ■ | ■ | ■ | ■ | ■ | ■ | + |
| altORF L6A | ■ | A | ■ | ■ | ■ | ■ | ■ | ■ | ■ | ■ | — |
| altORF L6K | ■ | K | ■ | ■ | ■ | ■ | ■ | ■ | ■ | ■ | — |
| altORF L6D | ■ | D | ■ | ■ | ■ | ■ | ■ | ■ | ■ | ■ | — |
| altORF W7A | ■ | ■ | A | ■ | ■ | ■ | ■ | ■ | ■ | ■ | + |
| altORF W7K | ■ | ■ | K | ■ | ■ | ■ | ■ | ■ | ■ | ■ | + |
| altORF W7D | ■ | ■ | D | ■ | ■ | ■ | ■ | ■ | ■ | ■ | — |
| altORF R8A | ■ | ■ | ■ | A | ■ | ■ | ■ | ■ | ■ | ■ | + |
| altORF R8L | ■ | ■ | ■ | L | ■ | ■ | ■ | ■ | ■ | ■ | + |
| altORF R8D | ■ | ■ | ■ | D | ■ | ■ | ■ | ■ | ■ | ■ | + |
| altORF I9A | ■ | ■ | ■ | ■ | A | ■ | ■ | ■ | ■ | ■ | + |
| altORF I9K | ■ | ■ | ■ | ■ | K | ■ | ■ | ■ | ■ | ■ | — |
| altORF I9D | ■ | ■ | ■ | ■ | D | ■ | ■ | ■ | ■ | ■ | — |
| altORF F10A | ■ | ■ | ■ | ■ | ■ | A | ■ | ■ | ■ | ■ | — |
| altORF C11A | ■ | ■ | ■ | ■ | ■ | ■ | A | ■ | ■ | ■ | — |
| altORF F12A | ■ | ■ | ■ | ■ | ■ | ■ | ■ | A | ■ | ■ | + |
| altORF R13A | ■ | ■ | ■ | ■ | ■ | ■ | ■ | ■ | A | ■ | — |
| altORF R13L | ■ | ■ | ■ | ■ | ■ | ■ | ■ | ■ | L | ■ | + |
| altORF R13D | ■ | ■ | ■ | ■ | ■ | ■ | ■ | ■ | D | ■ | + |
| altORF K14A | ■ | ■ | ■ | ■ | ■ | ■ | ■ | ■ | ■ | A | + |
| altORF K14L | ■ | ■ | ■ | ■ | ■ | ■ | ■ | ■ | ■ | L | + |
| altORF K14D | ■ | ■ | ■ | ■ | ■ | ■ | ■ | ■ | ■ | D | + |

**Proposed Consensus:** -X-L-{D/N}-X-{R/K/D/N}-$\Phi$-C-$\Phi$-(R/K)-X
